# Matrix maturation and cytoskeletal tension define strain thresholds for stretch-induced calcium signaling in human tendon cells

**DOI:** 10.1101/2025.04.10.648154

**Authors:** Patrick K. Jaeger, Fabian S. Passini, Barbara Niederoest, Maja Bollhalder, Sandro Fucentese, Jess G. Snedeker

**Author notes:** Correspondence to: Jess Snedeker, Lengghalde 5, 8008 Zurich, Switzerland.

## Abstract

The extracellular matrix (ECM) and mechanical loading shape cellular behavior, yet their interaction remains obscure. We developed a dynamic proto-tissue model using human tendon cells and live-cell calcium imaging to study how ECM and cell mechanics regulate mechanotransduction. Stretch-induced calcium signaling served as a functional readout. We discovered that ascorbic acid-dependent ECM deposition is essential for proto-tissue maturation and stretch-induced calcium signaling at physiological strains. Proto-tissue maturation enhanced stretch sensitivity, reducing the strain needed to trigger a calcium response from ∼40% in isolated cells to ∼5% in matured proto-tissues. A strong correlation between tissue rupture and calcium signaling suggests a mechanistic link to ECM damage. Disrupting ECM integrity, cell alignment, or cytoskeletal tension reduced mechanosensitivity, showcasing the influence of ECM and cytoskeletal mechanics on stretch-induced calcium signaling. Fundamentally, our work replicates calcium signaling observed in rodent tendon explants in vitro and bridges the gap between cell-scale and tissue-scale mechanotransduction.

**Teaser:** Matrix matters: tendon cells tune their response to stretch as their mechanical environment develops.

## Introduction

Tendon injuries and tendinopathy are common, worldwide public health problems. They are slow to heal and become chronic in up to a quarter of patients (*1*). They substantially reduce quality of life, decrease work productivity, and accelerate the development of diseases linked to a sedentary lifestyle (*2*), and for professional athletes, they can be career-ending (*3*). Healthy tendons consist of highly aligned collagen interspersed with relatively few cells (*4*). However, when injured, the cell behavior and the extracellular matrix (ECM) become dysregulated. There is increased cell proliferation, decreased cell alignment, and the collagen becomes disorganized and discontinuous (*5*). These changes weaken the tendon and make it more susceptible to further injury.

Physical therapy is currently the only established therapy for tendon injuries, and it is a lengthy process. Recovery takes weeks to months, and there is ongoing debate about the most effective loading regimen (*6*). Alternative therapies aim to promote tissue remodeling, but their efficacy remains highly controversial (*7*). Physical therapy, which primarily involves controlled mechanical loading, remains the gold standard approach and essential for recovery. Ongoing research is trying to elucidate the underlying mechanisms to enhance its efficacy with refined loading protocols or pharmacological agents. Studies have found that mechanical loading is not only important for recovery but also for development, homeostasis, and adaptation (*8*). While several sensors (e.g. focal adhesions, GPCRs, ion channels) and pathways (e.g. HIPPO, MAPK/ERK, Rho/ROCK) involved in mechanotransduction have been discovered (*9*), it is still unclear how and when these mechanisms regulate tendon biology, as the findings from in vitro studies have been inconsistent (*10*).

Drawing conclusions from in vitro studies that investigate mechanical loading is often challenging because they face limitations from artificial environments that neglect the extracellular matrix (ECM), and because they employ arbitrary loading protocols that are not grounded by physiological markers (*10, 11*). We previously identified one potentially important physiological marker using rodent tendon explants: we demonstrated that calcium signaling is a biochemical pathway that is activated upon mechanical loading of the tissue (*12*). Calcium signaling is a highly complex and versatile intracellular signaling system that controls various cellular processes (*13*). In tendon, we found that its regulation by mechanically activated ion channels guides tissue adaptation to extrinsic loading, and leads to improved elastic energy storage, which in turn translates to improved athletic performance (*12*).

While tendon explant models preserve the natural environment of tendon cells, they present significant challenges for studying the molecular mechanisms of mechanotransduction. These challenges include numerous limitations such as tissue degradation, difficulty in manipulating the ECM or cells, and difficulties implementing common biological assays due to the low cell density and high collagen content of tendons. Here, we address these limitations with a reductionist, in vitro approach that reproduces calcium signaling under physiological levels of strain and used it to investigate how ECM and cell mechanics influence mechanotransduction, focusing on calcium signaling as a physiological marker for functional mechanotransduction.

## Results

### Physiological strains trigger a calcium response in human tendon proto-tissues

Tendon cells experience stretch and shear stress when the tendon is mechanically loaded. To study stretch-activated mechanotransduction in greater detail, we built a uniaxial stretcher that is compatible with an inverted fluorescence microscope (Suppl. Fig. 1). We fabricated silicone chambers with a volume comparable to a single well of a 96-multi-well plate from 3D-printed molds and coated them covalently with collagen I. We cultured human tendon cells inside these chambers until they formed tendon proto-tissues, before stretching with concurrent, continuous live cell imaging (Fig. 1a). Using this model, we reproduced the stretch-induced calcium response previously observed in rodent tendon explants at physiological levels of strain (Fig. 1b; Suppl. video 1). Additionally, we implemented image analysis tools to track calcium signals in individual cells (Fig. 1c; Suppl. video 2).

**Fig. 1.**
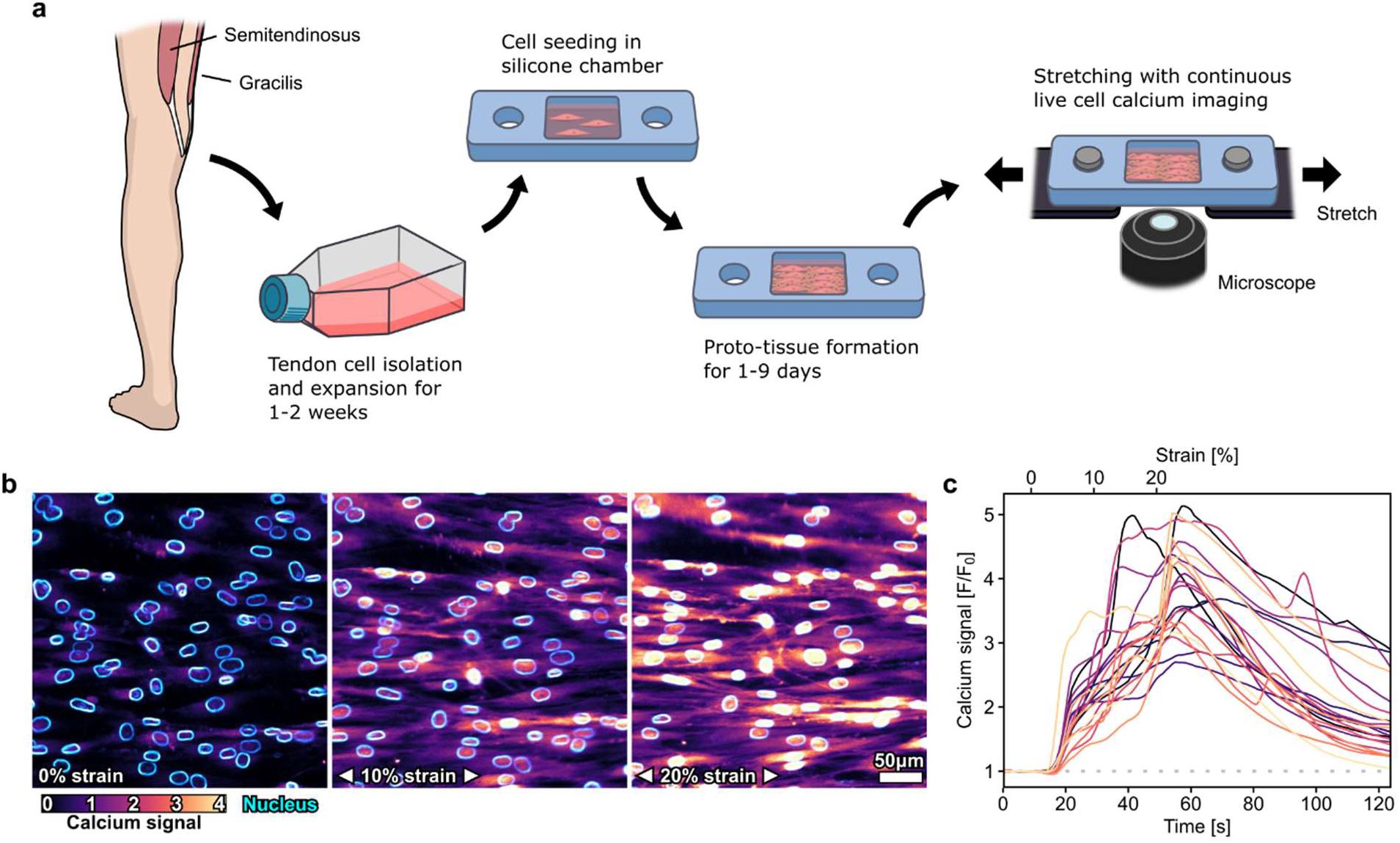
Continuous live cell calcium imaging of human tendon cells with concurrent stretching. (a) Schematic representation of the general experimental protocol. (b) Representative images showing the calcium response of tendon cells in a 7-day old tendon proto-tissue at rest and when stretched by 10% or 20%. Cells were stained with the fluorescent calcium reporter Fluo-4 AM and the DNA probe NucBlue. (c) Example of calcium signals of 22 cells shown in panel b while being stretched from 0-20% at a strain rate of 0.5%/s. The calcium response of individual cells is tracked in 250 ms intervals by averaging the Fluo-4 AM signal across the area of the nucleus.

### Cell-derived ECM deposition is critical for stretch-to-calcium mechanotransduction at physiological strains

In their native environment, tendon cells are embedded in a dense, collagenous ECM, which is the main load-bearing component of tendons. Therefore, we hypothesized that the ECM plays a crucial role in transmitting external forces to tendon cells. To test this hypothesis, we grew tendon proto-tissues in silicone chambers with or without ascorbic acid, an essential cofactor for collagen synthesis (*14*). When we stretched these proto-tissues, we found that without ascorbic acid-mediated ECM deposition the cells were drastically less sensitive to mechanical strain (Fig. 2a, b; Suppl. video 3). As expected, collagen deposition was almost absent without ascorbic acid after 3 days of culture (Fig. 2c), although the cell cytoskeleton was both prominently formed and anisotropically aligned to the loading axis of the substrate. Without ascorbic acid, the activation strain for calcium signaling increased by a factor of 2.9, and the calcium signal amplitude decreased by 40% (Fig. 2d-f). Thus, tendon cells failed to react to physiological levels of strain with a calcium response, in the absence of cell-derived ECM.

**Fig. 2.**
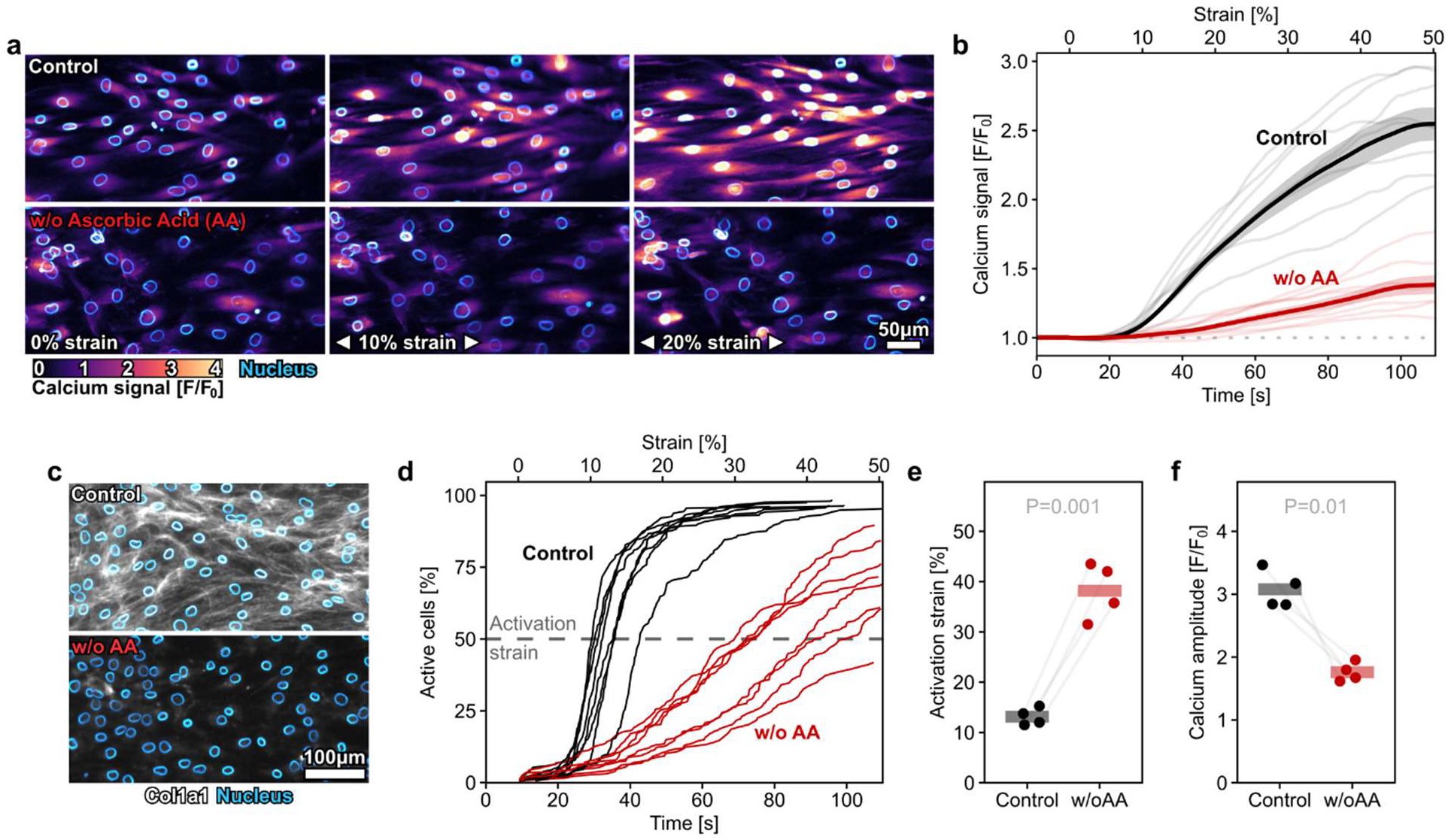
Ascorbic acid dependent extracellular matrix deposition is required for in vitro calcium signaling at physiological strains. (a) Example of calcium response of tendon cells stretched to 20% strain after 3 days in culture with and without supplementation of ascorbic acid. Data shown as mean±SE. (b) The average calcium response of cells cultured without ascorbic acid was substantially lower than that of the control group. (c) Tendon cells cultured for 3 days without ascorbic acid deposited substantially less collagen-I than the control group. Immunofluorescence staining against Col1a1. Nuclei were stained with NucBlue. (d) Cumulative activation curves of cells in tendon proto-tissues (one line represents one sample) cultured with and without ascorbic acid. Cells were classified as active once their calcium signal increased by more than 20% over the baseline. The activation strain of a proto-tissues was defined as the strain at which 50% of cells were classified as active (dashed line). (e) The calcium response of cells in tendon proto-tissues was triggered at a substantially higher activation strain when they were cultured without AA (+25.1±2.1% strain, mean±SE). (f) The amplitude of the calcium response was substantially weaker in cells cultured without AA (−1.3±0.2 F/F_0_, mean±SE). Tendon proto-tissues were cultured for 3 days and stretched to 50% strain at a strain rate of 0.5%/s. Data points represent biologically independent donors. Horizontal bars represent means. Statistical analyses were performed using linear mixed models with random intercepts for donors.

### Progressive proto-tissue formation sensitizes cells to physiological strains

The dependence on cell-derived ECM suggested that mechanotransduction relies on ECM deposition and remodeling, which requires time. We therefore continued to investigate how the duration of proto-tissue formation influences the calcium response. We observed that the activation strain stabilized between 4-5% strain after 5 days (Fig. 3a). Due to the viscoelastic nature of living cells and biological tissues, we also tested whether the strain rate influenced the tendon cells’ calcium response. We found that a very slow strain rate of 0.1%/s increased the activation strain and decreased the calcium signal amplitude (Fig. 3b-c), which is consistent with strain rate-dependent energy dissipation of viscoelastic materials. This effect was smaller in proto-tissues that were cultured longer before testing. Since we also observed considerable cell proliferation over time in the proto-tissues, we tested the influence of cell density on the cells’ sensitivity to strain (Suppl. Fig. 2). To do so, we adjusted the seeding density to grow proto-tissues with varying cell densities, independent of culture duration (Fig. 3d). We found no correlation between cell density and the calcium response. The cells’ calcium response was only reduced at very low (108±11 cells/mm^2^) and very high (951±90 cells/mm^2^) cell densities (Fig. 3e). These findings suggest that the mechanosensitivity of tendon cells is influenced more by the maturation and viscoelastic properties of the proto-tissue than by cell density alone.

**Fig. 3.**
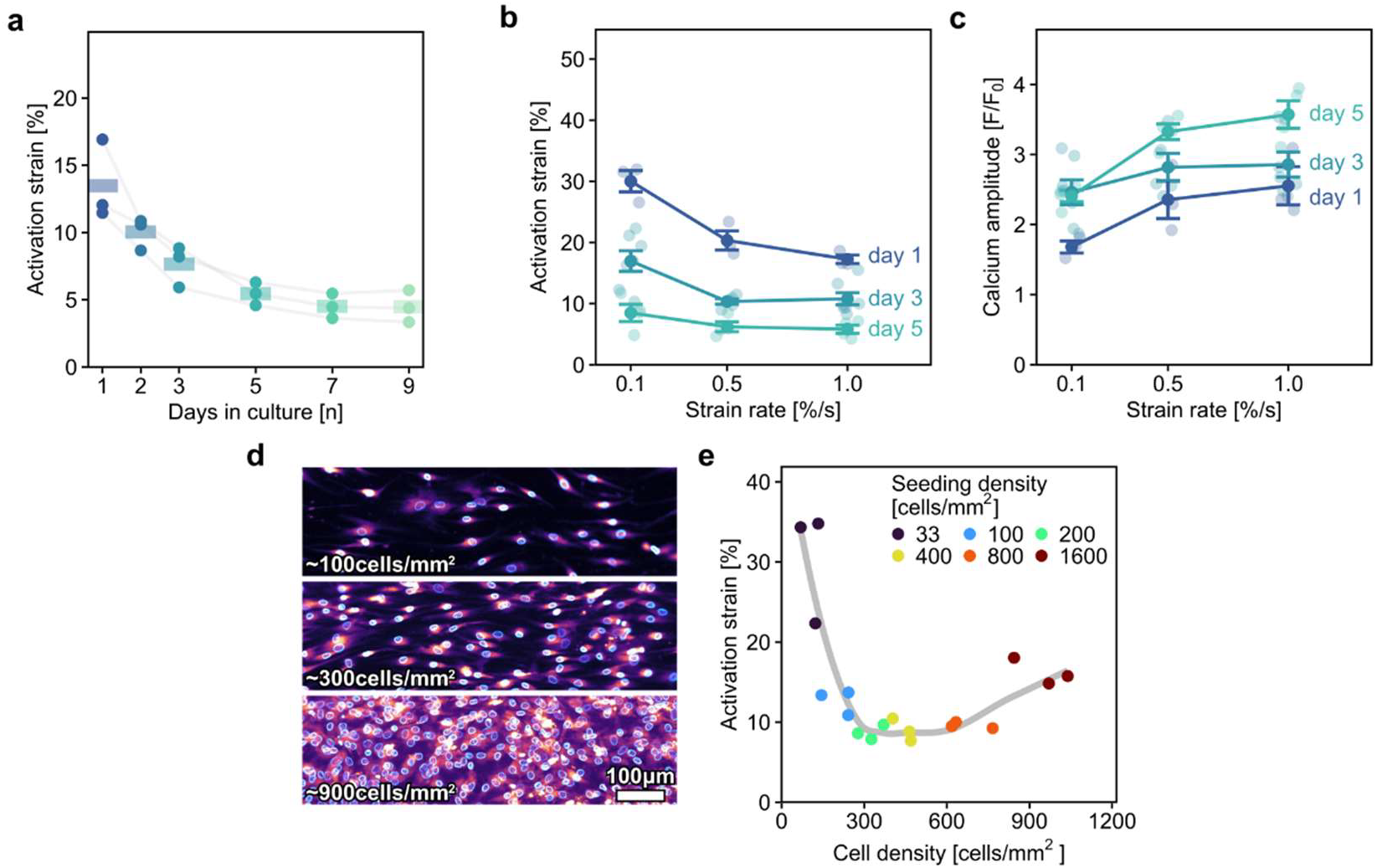
Progressive proto-tissue formation sensitizes the calcium response of cells to strains in the physiological range. (a) The calcium response of cells in tendon proto-tissues was triggered at progressively lower activation strains with prolonged culture duration and stabilized after 5-7 days. Stretching at a very low strain rate of 0.1%/s increased the activation strain. The effect was smaller in proto-tissues that were cultured longer before testing. Increasing the strain rate from 0.5%/s to 1.0%/s had a negligible effect on the activation strain. (c) Stretching at a very low strain rate of 0.1%/s decreased the amplitude of the calcium response. Increasing the strain rate from 0.5%/s to 1%/s only marginally increased the amplitude of the calcium response. (d) Examples of cell density after 3 days of culture with different seeding densities. (e) The activation strain did not correlate with cell density in proto-tissues that were seeded at different seeding densities and cultured for 3 days. Only very low and high seeding densities (33 and 1600 cells/mm^2^) increased the activation strain. Tendon proto-tissues were stretched at a strain rate of 0.5%/s to failure unless otherwise specified. Correlation was assessed using Pearson’s correlation coefficient.

### Disruption of ECM mechanics or integrity impedes stretch-induced calcium signaling

We observed that the failure strain (strain at onset of tissue rupture) of the proto-tissues decreased by 70% with longer culture durations, indicating that the tissue stiffness had increased over time (Fig. 4a-b; Suppl. video 4). We also detected a strong correlation between the failure strain and the activation strain (Fig. 4c). These findings indicated that there was not only increased ECM deposition over time but also progressive changes in ECM mechanics. We thus hypothesized that tissue stiffening accelerates the buildup of tension in the tissue when loaded, which ultimately triggered the calcium response at lower strains. To test this hypothesis, we softened the ECM by inhibiting ECM crosslinking with the lysyl oxidase inhibitor β-aminopropionitrile (BAPN). We indeed found that culturing the proto-tissues with BAPN increased the failure strain (Fig. 4d) and decreased the calcium response to mechanical loading (Fig. 4e-f). To further validate that tension in the ECM triggers calcium signaling, we treated proto-tissues with collagenase before stretching to disrupt the collagen network. Similarly, we observed an increased activation strain after collagenase treatment (Fig. 4g). These findings demonstrate that ECM stiffening enhances mechanotransduction by facilitating tension buildup, whereas disrupting ECM integrity impairs stretch-induced calcium signaling.

**Fig. 4.**
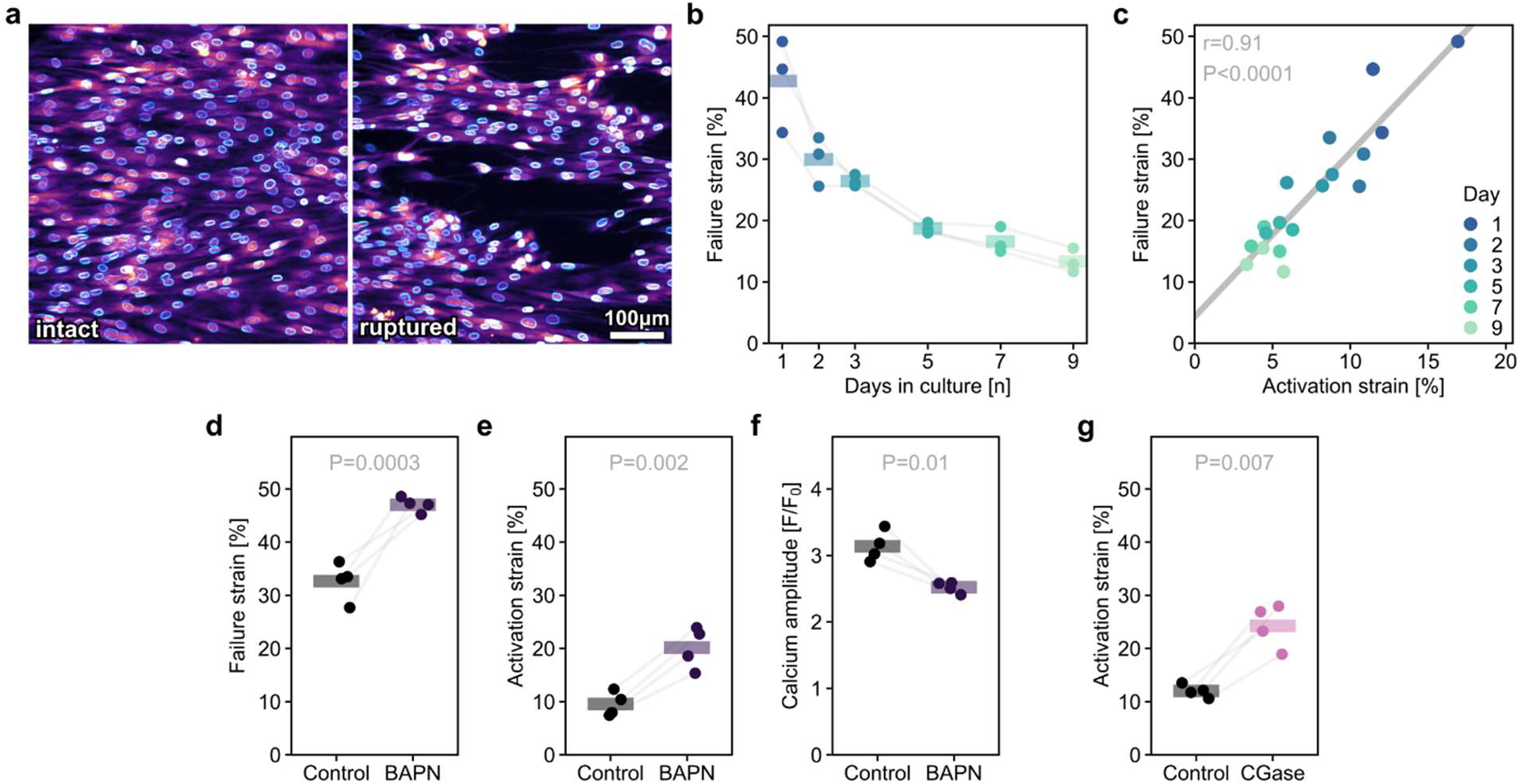
Extracellular matrix tension is required for the activation of stretch-induced calcium signaling. (a) Example of a tendon proto-tissue before and after rupture. (b) The failure strain (strain at onset of tissue rupture) decreased substantially over 9 days of culture (day 1 to day 9: -29.4±2.4% strain, mean±SE). (c) The activation strain correlated strongly with the failure strain in proto-tissues that were tested between 1-9 days of culture. (d) The failure strain of proto-tissues increased substantially (+14.3±1.9% strain, mean±SE) when matrix crosslinking was inhibited during the formation of proto-tissues with the lysyl oxidase blocker BAPN. (e) A higher activation strain (+10.6±1.0% strain, mean±SE) was required to trigger a calcium response in proto-tissues cultured with BAPN. (f) The amplitude of the calcium response was weaker (−0.6±0.1 F/ F_0_, mean±SE) in cells from proto-tissues that were cultured with BAPN. (g) Disruption of extracellular matrix integrity, with 20 minutes of collagenase treatment before stretching, increased the activation strain required to trigger a calcium response (+12.3±1.9% strain, mean±SE). Tendon proto-tissues were cultured for 3 days and stretched at a strain rate of 0.5%/s unless otherwise specified. Data points represent biologically independent donors. Horizontal bars represent means. Correlation was assessed using Pearson’s correlation coefficient. Statistical analyses were performed using linear mixed models with random intercepts for donors.

### Cytoskeletal tension is required for triggering stretch-induced calcium signaling

ECM tension translates to cytoskeleton (CSK) tension via focal adhesions and could thus activate ion channels that form complexes with focal adhesions or intracellular calcium stores. We therefore continued to investigate how CSK tension contributes to the stretch-induced calcium response by manipulating cell alignment and interfering with cytoskeleton polymerization before stretching. We manipulated cell alignment by rotating the substrate patterning (Fig. 5a) by 90°, shifting the major alignment axis of the cells to be perpendicular to the loading axis (Fig. 5a-b; Suppl. video 5). Perpendicular cell alignment increased the activation strain by 2.8-fold and decreased the amplitude of the calcium signals in active cells by one third (Fig. 5c-d). Next, we treated cells with Blebbistatin (to inhibit CSK pretensioning via myosin-II) or Cytochalasin D (to depolymerize actin) before stretching. These treatments increased the activation strain by 1.5- and 2.7-fold, respectively (Fig. 5e). These findings indicate that both, tension in the ECM as well as the CSK, are required for the proper activation of the relevant calcium channels for the mechanotransduction of mechanical strain into a calcium signal.

**Fig. 5.**
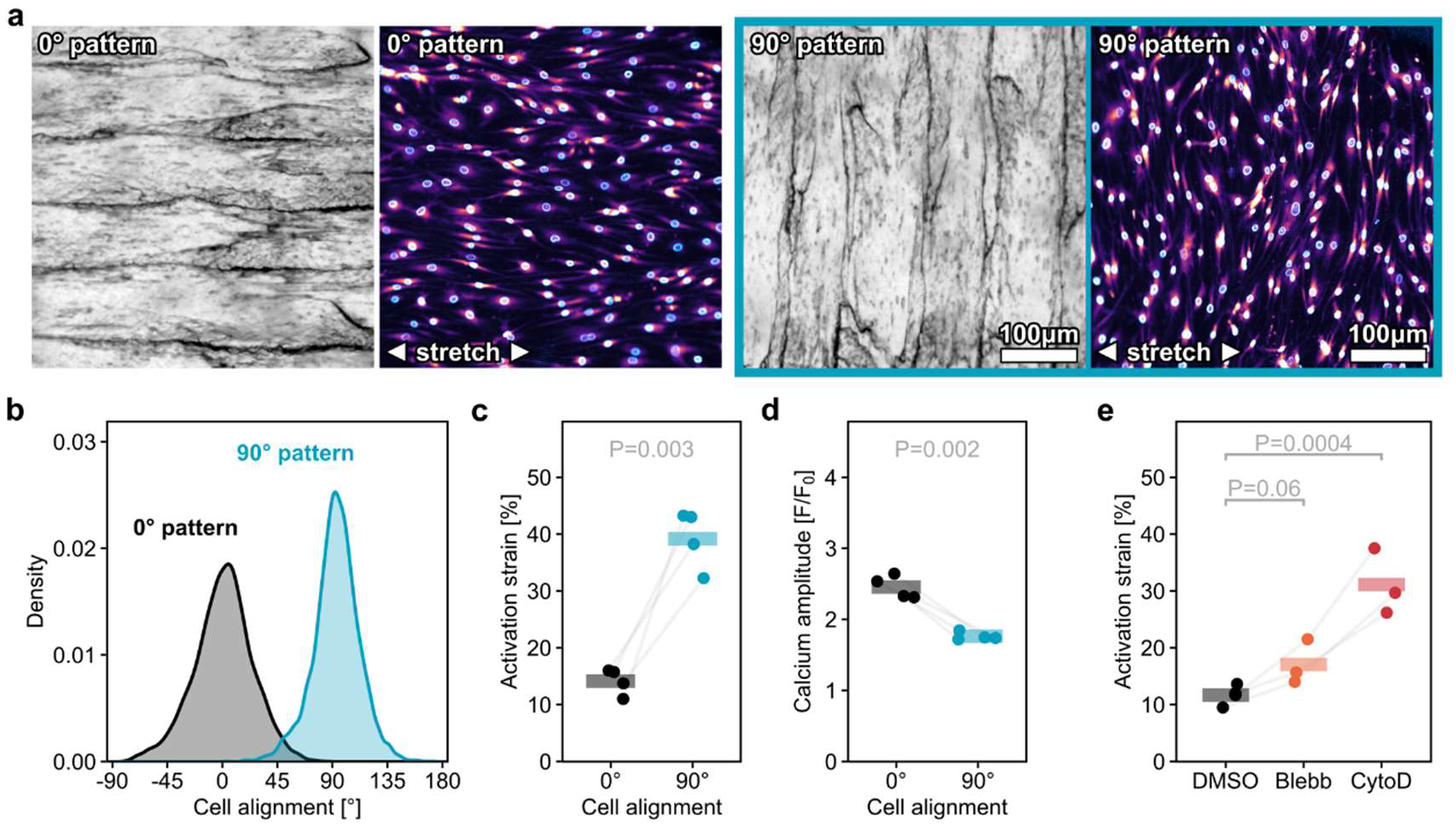
Cytoskeletal tension is required for the activation of stretch induced calcium signaling. (a) Example images of surface anisotropy created by casting silicone chambers from 3D printed molds, and resultant cell alignment. The direction of the surface anisotropy follows the nozzle movement of the FDM 3D printer. Cells aligned with the direction of the surface anisotropy. (b) Cells seeded in silicone chambers showed good alignment with the underlying surface anisotropy. Alignment quantification with ImageJ OrientationJ. A total of 16 samples from 4 biologically independent donors were analyzed. (c) A substantially higher activation strain (+25.1±2.9% strain, mean±SE) was required to trigger a calcium response in cells that were aligned at a 90° angle to the primary loading axis. (d) The amplitude of the calcium response was weaker (−0.7±0.06 F/F_0_, mean±SE) when the cells were aligned at a 90° angle to the primary loading axis. (e) Releasing cytoskeleton tension with Blebbistatin showed a trend towards increasing the activation strain (+5.6±2.3% strain, mean±SE), and depolymerizing the cytoskeleton using Cytochalasin D substantially increased the activation strain (+19.7±2.3% strain, mean±SE) required to trigger a calcium response. Data points represent biologically independent donors. Horizontal bars represent means. Statistical analyses were performed using linear mixed models with random intercepts for donors.

## Discussion

The molecular mechanisms behind mechanotransduction in healing and injured tendons remain largely elusive. In a previous study, we identified calcium signaling as a response to tendon overloading and found it to be involved in tendon adaptation (*12*). In the present study, we introduce an in vitro model that replicates this stretch-induced calcium response at physiological levels of strain. With this model, we demonstrate that ECM and cell mechanics interact, with a profound influence on calcium-mediated mechanotransduction.

Mechanotransduction in a tissue environment differs substantially from that in isolated cells. When testing proto-tissues grown with or without ascorbic acid, an essential cofactor for collagen synthesis and promotor of extracellular matrix (ECM) deposition (*15*), we observed that ECM deposition was crucial for replicating stretch-induced calcium signaling at physiological levels of strain. There could be several reasons that explain this change. Depending on the available binding motifs, different integrin types are recruited, which influence cytoskeletal dynamics and mechanical signaling cascades (*16, 17*). It has also been shown that focal adhesion formation in fibrous matrices, that allow remodeling by cells, is markedly different from that in rigid matrices or on flat substrates (*18*). Furthermore, it has been shown that cytoskeleton formation and tension beyond a basic ventral layer is necessary to trigger a strong, immediate calcium signal in response to mechanical actuation (*19*). Thus, the presence of cell-derived ECM could both alter cytoskeletal composition and structure. While single-cell models have helped deconstructing subcellular mechanisms, our findings suggest that tissue-level models are essential for accurately modeling mechanobiological responses relevant to tissues.

Several days of tissue formation were required to stabilize calcium-mediated mechanotransduction. We observed that over five days, the activation strain for triggering a calcium response dropped from close to 40% strain in isolated cells to 4.5% strain in matured proto-tissues, coming very close to the ca. 3%, observed in adult rodent tendon explants (*12*). We also observed that the strain rate-dependent effects on signaling dynamics decreased with prolonged culture. This change might be attributed to progressive inter-molecular bonding in the ECM that decreases the viscoelastic behavior of collagen matrices (*20, 21*).

ECM mechanics and integrity significantly influenced mechanotransduction. We observed that the failure strain approached an equilibrium after five days of culture at a level of ca. 13% strain, which is in the physiological range (*12, 22, 23*). Furthermore, the activation strain and failure strain were proportional, with tissue rupture occurring at 3-4 times the strain that initiated the calcium response. This ratio likewise matched the activation strain to failure strain ratio in rodent tendon (*12*). The decreasing failure strain with prolonged tissue culture indicates that proto-tissues stiffened over time. To test whether ECM mechanics influenced mechanotransduction, we altered the ECM stiffness through the inhibition of crosslinking enzymes. Consequently, the strain required to induce a calcium response increased in less crosslinked proto-tissues. Inducing micro-damage with collagenase had a similar effect. Altered tissue mechanics and ECM damage are both hallmarks of tendon disease (*24*). However, based on our findings alone it is difficult to draw conclusions on the precise impact of altered tissue mechanics on mechanotransduction after injury or in disease. Nonetheless, our results emphasize that ECM mechanics and damage must be controlled for dynamic in vitro studies to load cells reproducibly. Different cell lines or culture conditions can influence ECM composition and mechanics, and prolonged loading protocols or overloading can potentially damage the matrix and thus impair mechanotransduction. Based on these results, we propose that both the activation strain for calcium signaling, and the failure strain are relevant physiological markers that could be used to functionally anchor loading protocols in vitro.

Cytoskeleton (CSK) tension gates stretch-induced calcium signaling. We manipulated cells to align perpendicular to the loading axis and observed that the misaligned cells were substantially less sensitive to mechanical loading. We further confirmed that CSK tension is essential for initiating stretch-induced calcium signaling, as its pharmacological relaxation or depolymerization impaired the calcium response.Mechanically activated calcium channels generally fall into two main classes: those activated by tension in the lipid bilayer and those activated by tension transmitted from a tether (to the CSK or ECM), or both (*25*). Our results suggest that the channels that initiate the stretch-induced calcium response seem to be activated by tether-transmitted forces, since CSK tension was critical for initiating a calcium response. As for clinical relevance, changes in cell morphology and alignment are also hallmarks of tendon disease (*5*). However, it is difficult to say whether this is a cause of, or rather an adaptation to disease. Nonetheless, our results would suggest that cells in damaged or diseased tendons are potentially less responsive to mechanical loading. This supposition might explain why eccentric or higher-load physical therapy protocols seem to be most effective for the rehabilitation of tendon injuries (*26, 27*).

In conclusion, this study presents an in vitro model that closely reproduces physiological stretch-induced calcium signaling and emphasizes the critical roles of ECM mechanics and CSK tension in mechanotransduction. Moreover, we exploited it to highlight the distinction between tissue-scale and cell-scale mechanotransduction. These findings provide insight into important aspects of physiotherapy in humans and should help enhance the physiological relevance of dynamically stretched in vitro models - ultimately deepening our understanding of connective tissue regeneration and adaptation.

## Methods

### Silicone chamber fabrication

The molds for casting silicone chambers were 3D printed, with a Prusa MK3S equipped with a 0.25mm nozzle. Sylgard 184 silicone elastomer was used for casting and cured at 40°C for 24 hours. After demolding, the chambers were treated twice with sulpho-SANPAH (0.5mg/mL), dissolved in 50mM HEPES, for 10min in a Stratagene UV Stratalinker 2400. Then they were washed three times with deionized water and coated with collagen-1 (50μg/mL) from rat tails, dissolved in PBS, at 37°C for 3 hours. Finally, they were washed once with deionized water and stored at 4°C until use.

### Microscopy setup

The stretcher was constructed with two Zaber T-LSM025A linear stages (Zaber Technologies, Canada), 10×10mm aluminum extrusions and custom 3D-printed parts. The control software was programmed in Python. Images were acquired in rapid succession at 390nm and 488nm with 10ms and 50ms illumination time, respectively, with an inverted florescence microscope (Till photonics, Germany) at 10x magnification and a frame rate of 4 frames per second.

### Image analysis

The calcium signals of individual cells were analyzed using Fiji and the TrackMate plugin (Tinevez 2017). Individual cell nuclei were tracked based on images acquired at 390nm and the cells’ calcium signal was calculated by averaging the fluorescence signal of the corresponding image, acquired at 488nm, in a 7.5μm radius surrounding the centroid of the nucleus. Custom R scripts were used for further analyses. Cells were categorized as active once they showed an increase of 20% in calcium signal intensity over the baseline.

### Cell culture

Human tendon cells were isolated with collagenase from surgical debris of gracilis or semitendinosus tendons used during ACL-reconstructions at University Hospital Balgrist (with permission of Canton Zurich’s institutional review board and after informed consent from the patients). Cells were expanded on tissue culture plastic with DMEM (31885-023, Thermo Fisher), supplemented with 10% FBS (Thermo Fisher). Cells were used for experiments between passage 2 and 6. For seeding cells in silicone chambers, they were detached from tissue culture flasks using TrypLE (Thermo Fisher), resuspended in fresh medium, and seeded in silicone chambers. They were then cultured for 1-9 days, depending on the experiment before testing. The medium was exchanged every other day. Unless otherwise specified, the medium was supplemented with 200μM L-Ascorbic Acid Phosphate Magnesium Salt n-Hydrate (FUJIFILM Wako Chemicals, USA). For stretching experiments, cells were stained with degassed modified Krebs–Henseleit solution (KHS) containing 126 mM NaCl, 3 mM KCl, 2 mM CaCl_2_, 2 mM MgSO_4_, 1.25 mM NaH_2_PO_4_, 26 mM NaHCO_3_ and 10 mM glucose (3% oxygen, 29°C) with 3 μL/mL fluo-4 AM (Thermo Fisher), 40μL/mL NucBLue (Thermo Fisher), and pluronic F-127 (0.02% v/v) for 90 minutes before imaging. The staining was conducted at 3% oxygen, 29°C, and 5% CO2 to reduce the degradation of fluo-4 AM.

### Immunostaining

Samples were incubated with medium containing anti-collagen-I (1:200, PA2140-2, Boster) for 4 hours before washing with PBS and fixing with 4% paraformaldehyde. The secondary antibody (1:200, A-21202, A-21206, Thermo Fisher) was applied in PBS with 3% BSA and NucBlue (1:25 (v/v), Thermo Fisher) for 1 hour before washing again with PBS.

### Cytoskeleton, ECM manipulation

To inhibit crosslinking, BAPN (104470050, Fisher Scientific) was added to the medium at a concentration of 200μM for the duration of culture in the silicone chambers. To degrade the ECM, proto-tissues were incubated with KHS plus collagenase at a concentration of 2mg/mL for 20min before stretching. To relax the cytoskeleton, proto-tissues were incubated with KHS plus Blebbistatin (B0560, Sigma Aldrich) at a concentration of 10μM for 20min before stretching. To depolymerize the cytoskeleton, proto-tissues were incubated with Cytochalasin D (250255, Sigma Aldrich) at a concentration of 10μM for 20min before stretching.

### Statistical analysis

The calcium signals of proto-tissues cultured with or without are shown as means ± SE, alongside individual signals. Otherwise, horizontal bars are used to represent means alongside individual datapoints. Statistical analyses were performed using linear mixed models with random intercepts for donors using the lmerTest package in R (*28*). Correlation was assessed using Pearson’s correlation coefficient in R.

## Funding

The authors gratefully acknowledge funding of this study from the Swiss National Science Foundation, grant number 185095 “Mechanically triggered calcium influx - Seeking the molecular limit switch of the tendon mechanostat” and grant number 10000132 “TendO2n: targeting hypoxia signaling to combat human tendon disease”.

## Author contributions

Conceptualization: PKJ, JGS

Methodology: PKJ, FSP

Investigation: PKJ, MB, BN

Visualization: PKJ

Resources: SF

Writing—original draft: PKJ

Writing—review & editing: PKJ, JGS, FSP

## Competing interests

The authors declare that they have no competing interests.

## Data and materials availability

The main data supporting the findings of this study are available within the paper and its Supplementary Information. The raw and analyzed datasets generated during the study are too large to be publicly shared, yet they are available from the corresponding author on reasonable request.

## Code availability

The software of the stretching device, as well as MATLAB, ImageJ and R codes, are all available from the corresponding author on request.

## Supplementary

**Suppl. Fig. 1.**
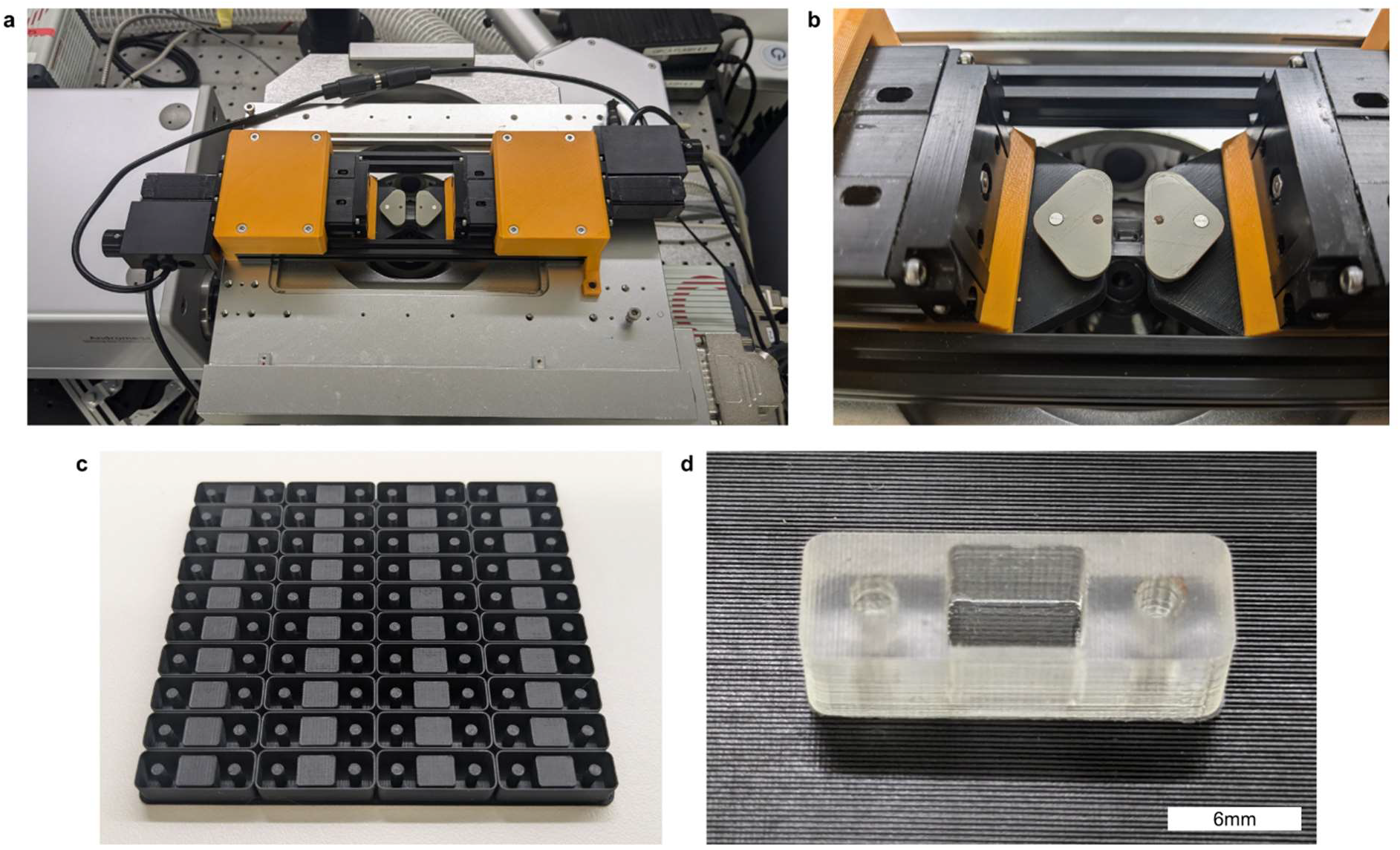
Hardware overview. (a) Stretcher mounted on the microscope stage. (b) Close-up of silicone chamber mounted in the stretcher. (c) 3D-printed mold for casting silicone chambers. (d) Close-up of silicone chamber.

**Suppl. Fig. 2.**
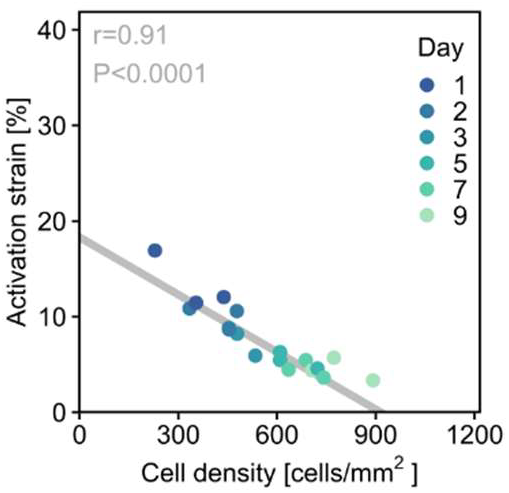
Correlation between culture duration and activation strain. The activation strain correlated strongly with the cell density in proto-tissues that were seeded at the same cell density (400 cells/mm^2^) and tested after 1-9 days of culture.

## Notes

### Competing Interest Statement

The authors have declared no competing interest.

